# Cdt1 modulates kinetochore-microtubule attachment stabilization via an Aurora B kinase-dependent mechanism

**DOI:** 10.1101/194993

**Authors:** Shivangi Agarwal, Kyle Paul Smith, Yizhuo Zhou, Aussie Suzuki, Richard J. McKenney, Dileep Varma

## Abstract

Robust kinetochore-microtubule (kMT) attachment is critical for accurate chromosome segregation. G2/M-specific depletion of human Cdt1 that localizes to kinetochores in an Ndc80 complex-dependent manner, leads to abnormal kMT attachments and mitotic arrest. This indicates an independent mitotic role for Cdt1 in addition to its prototypic function in DNA replication origin licensing. Here, we show that Cdt1 directly binds to microtubules (MTs). Endogenous or transiently expressed Cdt1 localizes to both mitotic spindle MTs and kinetochores. Deletion mapping of Cdt1 revealed that the regions comprising the middle and C-terminal winged-helix domains but lacking the N-terminal unstructured region was required for efficient MT-binding. Mitotic kinase Aurora B interacts with and phosphorylates Cdt1. Aurora B-phosphomimetic Cdt1 exhibited attenuated MT-binding and its cellular expression induced defective kMT attachments with a concomitant delay in mitotic progression. Thus we provide mechanistic insight into how Cdt1 affects overall kMT stability in an Aurora B kinase phosphorylation-dependent manner; which is envisioned to augment the MT-binding of the Ndc80 complex.

**eTOC summary:** • Cdt1 binds to microtubules

• The middle and the C-terminal winged-helix domains of Cdt1 are involved in MT-binding

• Aurora B Kinase phosphorylates Cdt1 and influences its MT-binding

• Aurora B-mediated Cdt1 phosphorylation is necessary for kMT stability and mitotic progression

## Introduction

Accurate chromosome segregation during mitosis is accomplished by the concerted function of the bipolar mitotic spindle and kinetochores (Westhorpe and Straight, 2013). In this process, mitotic cells inevitably confront a challenge to maintain robust kinetochore-microtubule (kMT) attachment despite dynamic instability generated by rapid polymerization (growth) and depolymerization (shrinkage) of microtubules (MTs) (Joglekar et al., 2010; DeLuca and Musacchio, 2012). How this dynamic process of kMT coupling is accomplished and regulated in vertebrates is not well understood.

A highly conserved network of protein complexes, called the KMN network (Knl1, Mis12, Ndc80) is the core interface that links spindle MTs to the kinetochores (Cheeseman et al., 2006; Cheeseman and Desai, 2008; Afreen and Varma, 2015). The N-terminal region of the Ndc80 complex, containing the Calponin homology (CH) domain and positively charged tail domain of the Hec1 subunit, constitute an essential MT-binding site (Varma and Salmon, 2012). The N-terminal region of Knl1 has also been shown to be necessary for MT binding (Cheeseman et al., 2006; Welburn et al., 2010; Espeut et al., 2012). Recently, an unprecedented mitotic role for human Cdt1, a well-established DNA replication licensing factor, was discovered (Varma et al., 2012; Pozo and Cook, 2016). The Hec1 loop domain that generates a flexible hinge in an otherwise rigid Ndc80 complex (Wang et al., 2008), has been shown to recruit Cdt1 to kinetochores by interacting with Cdt1’s N-terminal region (Varma et al., 2012). Precise high-resolution separation measurement (delta) analysis between the extreme N- and C-termini of Ndc80 revealed that the Ndc80 complex bound to Cdt1 maintains an extended conformation that serves to stabilize kMT attachments via an unknown mechanism (Varma et al., 2012). Thus, while substantial research has provided insights into the structural and mechanistic aspects of the “canonical” licensing function of Cdt1, how Cdt1 influences kMT attachments during mitosis remains unclear.

Besides Cdt1, the loop region of Hec1 also serves as a docking site for several other microtubule-associated proteins (MAPs) such as Dis1 (vertebrate homolog of chTOG) and Dam1 in yeast (Hsu and Toda, 2011; Maure et al., 2011; Schmidt and Cheeseman, 2011). In fact, in budding yeast, the Ndc80 and Dam1 complexes function synergistically to bind to MTs (Tien et al., 2010). Similarly, in vertebrates the loop region has been reported to recruit the Ska complex (Zhang et al., 2012) that also interacts with MTs through the unique winged-helix domain (WHD) of the Ska1 subunit. Further, the Ndc80 complex increases the affinity of the Ska1 subunit for MTs by 8-fold (Schmidt et al., 2012; Abad et al., 2014). These studies suggest that although the Ndc80 complex is critical for kMT-binding, other factors such as the Dam1 and Ska complexes are required to efficiently orchestrate kMT attachments and chromosome segregation.

The present study was thus undertaken to address critical outstanding questions surrounding the role of Cdt1 at kinetochores in stabilizing kMT attachments (Varma et al., 2012). These include: (1) whether Cdt1 directly binds to MTs, and (2) how Cdt1 is regulated for its mitotic function. Employing several biochemical, biophysical and cell biological approaches, we demonstrate that human Cdt1 can directly interact with MTs of the mitotic spindle. We further show that Cdt1 is a novel target for Aurora B Kinase and that Aurora B-mediated phosphorylation of Cdt1 regulates its MT-binding properties that in turn influences mitotic progression.

## Results

### Cdt1 directly binds to microtubules *in vitro*

We had previously demonstrated that Cdt1 localizes to mitotic kinetochores, dependent on the loop domain of the Hec1 subunit of the Ndc80 complex. Further, using a novel RNAi-mediated knockdown approach and microinjection of a function-blocking Cdt1 antibody, we showed that perturbation of Cdt1 function specifically during mitosis led to unstable kMT attachments culminating in a late prometaphase arrest (Varma et al., 2012). Moreover, high-resolution microscopic analysis suggested that in the absence of Cdt1, the coiled coil of the Ndc80 complex assumed a bent conformation and that the complex was not able to make a full extension along the kMT axis (Varma et al., 2012). But how Cdt1 contributed to this mechanism and imparted kMT stability was unclear.

To investigate this mechanism systematically, we began by analyzing the structure of Cdt1 but since no high-resolution structures of full-length Cdt1 have been published, we first subjected the human Cdt1 amino acid sequence to at least ten different secondary structure prediction algorithms that predict disordered regions. Although, the consensus readily identified previously crystallized winged helix domains (WH) as being well-folded; the N-terminal 93 amino acids were convincingly predicted to be at least ~35% intrinsically disordered, in addition to the “linker” region between the two WH domains **(Fig. 1 A)**. Homology models generated using three modeling programs, reliably identified the middle (WHM) and the C-terminal winged helix (WHC) domains, while the N-terminal region (1-160) was consistently predicted to have only limited secondary structure elements in the absence of other binding partners **(Fig. 1 B)**. The NMR solution structure (PdbID: 2KLO) determined for the C-terminal 420-557 aa of *Mus musculus,* abbreviated as mCdt1, revealed that this region adopts a WH domain conformation (Khayrutdinov et al., 2009) in addition to the other WH domain in the middle of Cdt1 (172-368 aa) responsible for Geminin binding (Lee et al., 2004). The WH domain has traditionally been implicated primarily as a DNA binding motif, but emerging evidences suggest that it could also serve as a protein-protein interaction domain (Khayrutdinov et al., 2009). In addition to the canonical three helix-bundles and anti-parallel beta-sheets, the Cdt1 WHC also contains an additional helix (H4) at the C-terminus (**Fig. S1 A**). Superimposition of the homology model for human Cdt1 fragment 410-546 aa [generated using Phyre2 (Kelley et al., 2015)] with the available NMR structure of an mCdt1 fragment showed an overlap with the root mean square deviation of 1.038 Å, indicating that the 410-546 aa of human Cdt1 indeed conforms to a WH domain (**Fig. 1 B**).

**Figure 1.**
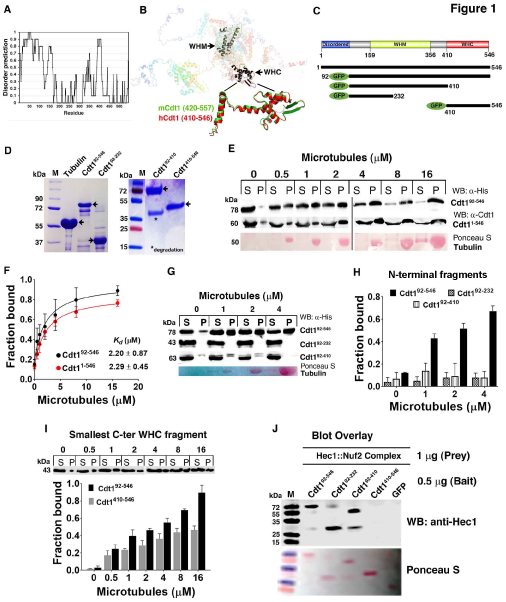
Cdt1 is a novel microtubule-binding protein. (A) Disorder Prediction in human Cdt1. Disordered prediction (sum of 10 programs) is plotted as a function of residue number. (B) Seven homology models of full-length Cdt1; colored from N-terminus (blue) to C-terminus (red) with regions of low confidence or low overlap shown in translucent. Conserved WHM and WHC domains shown and aligned in grey and black, respectively, for clarity. Magnified is the PyMol rendered cartoon representation showing superimposition of the C-terminal human Cdt1 (410-546 aa, in red) generated using Phyre2 server with available NMR structure (PDBid: 2KLO, in green) of mouse C-terminal Cdt1 (420-557 aa). (C) Constructs used in this study, full-length Cdt1 (1-546 aa), its deletion variant devoid of N-terminal 92 aa (92-546) and N- and C-terminal deletion variants generated in fusion with an N-terminal GFP tag for expression in bacteria are shown as black bars. Disordered domain shown in blue, winged helix middle (WHM) domain shown in yellow, and winged helix C-terminal (WHC) domain shown in red. (D) SDS-PAGE (15%) of the indicated purified recombinant proteins along with native tubulin purified from porcine brain. M, molecular mass standard. Asterisk (*) denotes degradation bands (confirmed by western blot) in Cdt1^92-410^. Arrows point towards the band corresponding to the protein of interest. (E) Western blots showing MT co-sedimentation for Cdt1^92-546^ (GFP-tagged) and full-length Cdt1^1-546^ (without any tag), 1 μM each; purified from bacteria with the indicated concentrations of taxol-stabilized MTs (in μM). Samples fractionated as Supernatant (S) and Pellet (P) were analyzed by western blot probed with antibodies against 6×-His tag or anti-Cdt1 as indicated in each case to detect Cdt1. Ponceau S stained tubulin. (F) Quantification (Mean ± SD, n=3) of the Cdt1^92-546^ and Cdt1^1-546^ blots using Image J. The 95% CI values obtained for the fit were 0.60-0.94 for *B_max_,* 0.37-4.04 for *K_d_* and 0.07-0.29 for the background for the former and 0.63-0.79 for *B_max_,* 1.33-3.25 for *K_d_* and 0.08-0.18 for the background for the latter construct. (G) Representative western blots of the binding of indicated deletion fragments to MTs in a co-sedimentation assay with the indicated MT concentrations (top). (H) Quantification of binding (Mean ± SD, n=3) of the deletion fragments to MTs. (I) Quantification of MT-binding of Cdt1’s smallest C-terminal fragment, Cdt1^410-546^ and a representative western blot (top) from three independent experiments (Mean ± SD, n=3). (J) Blot overlay assay to study Cdt1-Hec1 interaction. Indicated proteins (0.5 jg each) were loaded as baits on 18% SDS-PAG, transferred to nitrocellulose membrane and blocked with 5% SM-TBST. Hec1::Nuf2-His dimeric complex (1 |jg) was overlaid as prey protein on the membrane for 12 h at 4 °C. The blot was washed and probed with anti-Hec1 antibody (1:2000, Abcam 9G3) followed by chemiluminescence. The same blot was probed with Ponceau S.

The presence of a central and a C-terminal WH domain in human Cdt1 similar to the MT-binding domain of the Ska complex (Abad et al., 2014) prompted us to posit a new paradigm where Cdt1 bound to the loop may directly bind to MTs to create an additional MT-attachment site for the Ndc80 complex and contribute to the robustness of kMT attachments. To test our hypothesis, we first evaluated the ability of Cdt1 to directly bind to MTs using *in vitro* assays.

Proteins with large disordered regions are generally characterized by rapid degradation (Vavouri et al., 2009). Indeed, challenges in purifying the full-length human Cdt1 from bacterial system were circumvented by the deletion of the 92 aa disordered stretch from its N-terminus which likely contributed to its degradation, reduced expression and poor solubility. The Cdt1 protein was successfully purified with an N-terminal GFP fusion (abbreviated as Cdt1^92-546^, **Figs. 1, C and D**) and was evaluated for its ability to bind to MTs in a co-pelleting assay. The Cdt1^92-546^(1 μM) was found to co-pellet efficiently with taxol-stabilized MTs (**Fig. 1 E**). This interaction with MTs is specific as Cdt1 was not enriched in the pellet fraction in the absence of MTs. To determine the apparent MT-binding affinity of Cdt1^92-546^, we varied the MT concentration and measured the Cdt1 concentration required for half maximal binding. In this assay, the apparent dissociation constant (K_d_) was determined to be 2.20 ± 0.87 μM (**Fig. 1 F**). Since the construct used in this assay was devoid of the first 92 aa, full-length (FL) Cdt1^1-546^ was purified with a GST tag which was subsequently cleaved using Factor Xa and the untagged Cdt1^1-546^ was tested for its ability to bind to MTs. As expected, the full-length protein also bound to MTs (**Fig. 1 E**) with an affinity (2.29 ± 0.45 μM) similar to the Cdt1^92-546^ (**Fig. 1 F)**; indicating that the first disordered 92 aa are dispensable for the MT binding function of Cdt1. As controls, neither purified GFP nor GST proteins fractionated with MTs substantiating the specificity of interaction **(Fig. S1, B and C)**.

To map the Cdt1-MT interaction, deletion fragments within Cdt1^92-546^ were generated (**Figs. 1, C and D**). Interestingly, neither of the N-terminal fragments, Cdt1^92-232^ or Cdt1^92-410^ cosedimented with MTs (**Figs. 1, G and H**). However, the smallest C-terminal fragment Cdt1^410-546^ did show MT binding, albeit its efficacy was reduced by ~50% as compared to the Cdt1^92-546^ (**Fig. 1 I**). Thus, we conclude that the C-terminus of Cdt1 (410-546 aa) is necessary but not sufficient for MT binding.

We further tested the ability of Cdt1 deletions to bind to Hec1 in an *in vitro* blot overlay assay. In accordance with the previous study (Varma et al., 2012), our data shows that although the N-terminal fragments, Cdt1^92-232^ and Cdt1^92-410^ were incompetent to sediment MTs (**Fig. 1 G**); they interacted with Hec1 in this assay (**Fig. 1 J**). On the other hand, the Cdt1^410-546^ that did bind MTs to some extent was not able to bind Hec1 (**Fig. 1 J**). GFP loaded as control bait protein was unable to bind Hec1 prey.

### Cdt1 preferentially binds to straight MTs independent of the E-hook tails

At the C-terminus of tubulin are negatively charged acidic tails of ~20 aa, known as “E-hooks” that represent a critical recognition site for many MT binding proteins including Ndc80^Bonsai^ (Ciferri et al., 2008) and Dam1 (Ramey et al., 2011). The ability of Cdt1 to bind to MTs devoid of E-hooks was thus assessed in a co-sedimentation assay. Similar to Ska1 (Abad et al., 2014) and chTOG (Spittle et al., 2000), Cdt1^92-546^ did not show any appreciable reduction in its ability to bind to MTs lacking E-hooks (**Figs. 2, A and B**). This suggests that the critical contacts involved in this interaction might be the structured regions of tubulin monomers rather than the unstructured acidic tails. The highly basic nature of Cdt1 (predicted pI=9.9) is indicative of MT recognition through electrostatic/ionic interactions. Indeed, the binding of Cdt1^92-546^ with MTs was substantially reduced to ~50% in the presence of 250 mM NaCl (**Fig. 2 C**); suggesting that the binding may be at least partially electrostatic in nature.

**Figure 2.**
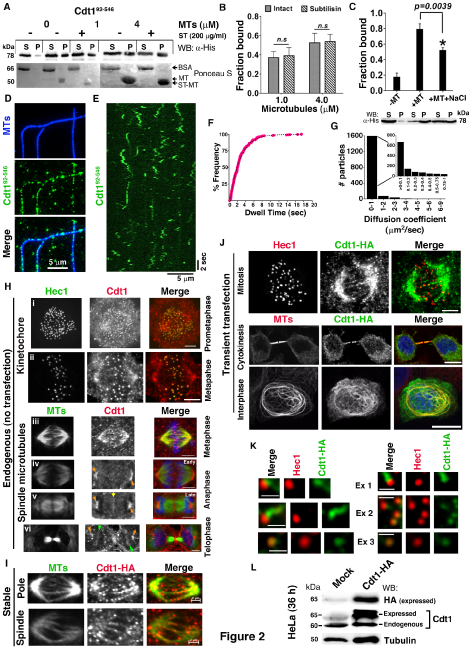
Cdt1 diffuses on MTs *in vitro* and decorate spindle microtubules *in vivo.* (A) Representative western blot of Cdt1^92-546^ (1 μM) co-sedimentation with intact (MT) or subtilisin-treated (ST-MT) microtubules at 1 and 4 μM concentrations. (B) Quantification of (A) (Mean ± SD, n=3). (C) Quantification of co-sedimentation of Cdt1^92-546^ (1 μM) with 16 μM MTs in the absence (+MT) and presence of 250 mM NaCl (+MT+S) (top). Representative western blot shown (bottom). (D) Single channel and merged TIR-FM images showing surface immobilized Dylight405-labeled MTs (blue) and GFP-tagged Cdt1^92-546^ (green). (E) Kymograph analysis of Cdt1^92-546^ interactions with the MT lattice. (F) Cumulative frequency plot and exponential fit of Cdt1^92-546^ dwell times. N=304 molecules, 2 independent protein preparations. (G) Diffusion coefficient analysis of Cdt1^92-546^ on MTs carried out using DiaTrack particle tracking Software. Inset graph shows the histogram distribution of number of particles of Cdt1^92-546^ that diffused on MTs within the range of 0-1 μm^2^/s. (H) Untransfected mitotic HeLa cell stained with anti-Cdt1 and either anti-Hec1 antibody (top panels i and ii) or anti-tubulin antibody (bottom panels iii-vi). Scale bar, 5 μm. Shown are the representative images for Cdt1 kinetochore (anti-Cdt1/red and anti-Hec1/green) and spindle (anti-Cdt1/red and anti-tubulin/green) staining at different stages of mitosis in HeLa cells as indicated. Orange arrows in panels iv-vi depict Cdt1 staining at the spindle poles; yellow arrow in panel v depicts the Cdt1 staining at the cleavage furrow in late anaphase and green arrows in panel vi depict Cdt1 localization to the reforming nucleus in telophase. Immunostaining of stably transfected (I) or transiently transfected (J) HeLa cells with anti-HA, anti-Hec1 and anti-tubulin antibodies; as indicated. Scale bar, 5 μm. The middle and bottom panels in (J) depict confocal images of cells in cytokinesis and interphase, respectively, with DAPI staining of chromosomes in blue. (K) Insets from the image stack of the mitotic cell in the top panels of J (left, also see Supplementary movie S2) with an example from another cell in Supplementary movie S3 (right). Scale bar, 1 μm. (L) Western blot of mock and transfected lysates to probe for Cdt1 (using anti-HA or anti-Cdt1 as indicated) and mouse tubulin as loading control. “Expressed” and “Endogenous” represent ectopically expressed (with C-terminal 2×-HA, S- and 12×-His-tags; abbreviated as HASH tags) and endogenous (without tag) Cdt1, respectively.

Further, we assessed the preference for Cdt1 to bind to straight *vs* curved MTs. Unlike the CH domains of Ndc80 that bind to the dimeric tubulin interface and prefer a straight MT protofilament conformation; Ska accesses MTs by interacting with the structured regions within tubulin monomers independent of any conformation (Abad et al., 2014). The data revealed that even though MT-binding was considerably reduced when the MTs were not straight; Cdt1 did retain the capacity to bind to curved MT protofilaments (**Fig. S1, D-F**), which may have important implications for the function of Cdt1 at the kMT interface in conjunction with the Ndc80 complex.

In order to test the preference of Cdt1 to bind to MT ends, polymerized MTs were sheared to increase the number of ends. We did not observe any significant increase in the association of Cdt1 with the MTs in this scenario (**Fig. S1, D-F**). This suggests that Cdt1 might not be merely a MT plus/minus end-specific protein, but its binding sites are likely distributed along the length throughout the MT lattice.

### Cdt1 diffuses on MTs *in vitro* and decorates mitotic spindles *in vivo*

To substantiate Cdt1-MT binding further, we directly visualized Cdt1^92-546^ interaction with MTs using total internal reflection fluorescence microscopy (TIR-FM). The Cdt1^92-546^ bound to Dylight405-labelled MTs at concentrations as low as 1 nM (**Fig. 2 D)** and at higher concentrations (10 nM or higher), it completely decorated the entire MT lattice **(Fig. S1 G)**. In our conditions, individual molecules either bound statically or diffused along the MT lattice **(Fig. 2 E)**. The dwell time distribution for Cdt1^92-546^ on MTs was well described by a single exponential decay function yielding a characteristic dwell time (τ) of 2.37 sec **(Fig. 2 F)**. The calculated association rate (K_on_,10 ^-6^,s ^-1^,nM ^-1^) was 1.07 ± 0.38 and the dissociation constant (K_D_, μM) was 0.46 ± 0.17 (**Fig. 2 F**). A large fraction of Cdt1^92-546^ (~65%) was found to undergo short-range diffusive motion along the MT lattice to a moderate extent that falls in the range of 0-0.5 μm^2^/sec **(Fig. 2 G, E and Movie S1)**. A smaller fraction (~20%) did not apparently diffuse at all while an even smaller fraction (~15%) exhibited relatively fast diffusion (>1 μm^2^/sec). The average diffusion coefficient for Cdt1^92-546^ was ~0.16 μm^2^/sec under our experimental conditions (**Fig. 2 G**). Both Ndc80 and Ska complexes have also been shown to exhibit diffusion along the MT lattice; albeit at variable rates comparable to that observed for Cdt1 under our assay conditions (Schmidt et al., 2012; Zaytsev et al., 2015).

Although the *in vitro* experiments provide evidence that Cdt1 binds to MTs, we sought further evidence for this activity in cells. We had previously observed Cdt1 localization at mitotic kinetochores in paraformaldehyde-fixed LLCPK1 and PTK2 cells (Varma et al., 2012). We now show Cdt1 localization to kinetochores in HeLa cells using identical fixation conditions (**Fig. 2 H**). While Cdt1 transiently localized to the kinetochores predominantly during late prometaphase and metaphase (**Fig. 2 H, panels i and ii**), the detection dropped to almost negligible levels in anaphase and telophase (not shown). Immunofluorescence staining of Cdt1 in methanol-fixed HeLa cells, which removed most of the kinetochore staining, allowed us to visualize clear mitotic spindle staining (**Fig. 2 H)**. The variability observed in our previous (Varma et al., 2012) and present studies with respect to Cdt1 localization to kinetochores or spindle MTs is attributed to different antibodies, fixation conditions and the type of cells used.

To evaluate the spindle localization of Cdt1 comprehensively, HeLa cells at different stages of mitosis were examined. Anti-Cdt1 antibody staining substantially overlapped with the mitotic spindle but individual spindle MT staining was often hard to discern. Cdt1 staining on spindles was intermittent with areas of brighter or dimmer intensities; with maximum intensity observed at the spindle poles. Cdt1 spindle staining started to appear in late prometaphase after majority of the chromosomes were aligned with no evident MT staining during early or mid-prometaphase (not shown). Cdt1 spindle staining was the brightest in metaphase (**Fig. 2 H, panel iii**). A low magnification image with several cells demonstrating similar Cdt1 co-localization with spindle MTs during metaphase is shown (**Fig. S1 H**). In anaphase and telophase, Cdt1 MT staining is predominantly restricted to the spindle poles as discrete crescent-shaped structures that are brighter in anaphase compared to telophase (**Fig. 2 H, panels iv-vi**). No discernable Cdt1 localization to inter-polar MTs or the MT bundles of the central spindle regions in anaphase or telophase was observed. Interestingly but unexplainable as of now, Cdt1 staining at the cell periphery became prominent at the beginning of anaphase and in telophase, with moderately bright staining appearing at the regressing cleavage furrow (**Fig. 2 H, panels v and vi**). Faint Cdt1 co-localization was also observed in the condensing nuclei in late telophase consistent with its G1 nuclear function (**Fig. 2 H, panel vi**).

To lend further support to these observations, full-length Cdt1 was transiently overexpressed with an HA tag in HeLa cells and immunostaining was performed to assess its localization. In agreement with the results so far, staining with anti-HA clearly reveals localization of Cdt1-HA on the spindle MTs and also on kinetochores (overlap with Hec1, a kinetochore marker required for Cdt1 recruitment) in mitotic cells (**Fig. 2 J, top panels; Movies S2 and S3**). As shown in the **Fig. 2 K**, magnification of selected kinetochores from multiple cells further validates Cdt1 localization to kinetochores and kMTs (**Fig. 2 K; Movies S2 and S3**). Strikingly, interphase cells expressing high levels of Cdt1 showed a MT bundling phenotype (**Fig. 2 J**). This was not evident in cells with low or moderate expression of Cdt1 which is similar to the previous findings of Welburn *et al*; where a bundling phenotype was observed only when GFP-Ska1 was over-expressed *in vivo* or when higher concentration of Ska complex was used *in vitro* (Welburn et al., 2009). Further, in cells that were undergoing cytokinesis, Cdt1 was localized to MT-bundles at the mid-body region (**Fig. 2 J**). Western blotting of transfected HeLa cell lysates with anti-HA (representing the exogenously expressed Cdt1 with HA tag) confirmed expression of Cdt1 as full-length protein in fusion with the HA tag (**Fig. 2 L**). Similar MT-staining (spindle/spindle pole) was also obtained in HeLa cells that stably expressed HA-tagged Cdt1, representing a scenario between over- and endogenous expression (**Fig. 2 I**).

### Aurora B Kinase phosphorylates Cdt1 and affects Cdt1-MT binding *in vitro*

We then sought to understand how the MT-binding of Cdt1 could be regulated. Aurora B phosphorylates multiple substrates at kinetochores to negatively regulate kMT attachments and is also instrumental in correcting erroneous attachments (Welburn et al., 2010; Chan et al., 2012; Schmidt et al., 2012). *In vitro* studies with purified Ndc80 complex demonstrated that introduction of either multiple phosphorylations or phosphomimetic substitutions within the Hec1 tail reduced Ndc80-MT binding, while a lack of phosphorylation promoted stronger binding (Cheeseman et al., 2006; Umbreit et al., 2012). Similarly, Ska1, another kinetochore-localized MAP is a substrate for Aurora B and its phosphorylation at four sites located within the MTBD drastically reduces its kinetochore localization (Welburn et al., 2010; Chan et al., 2012; Schmidt et al., 2012). Mutating two of these, S185 and S242 to aspartate dramatically reduced the MT binding activity of the Ska complex *in vitro* and resulted in a mitotic delay and reduced kMT stability *in vivo* (Chan et al., 2012; Schmidt et al., 2012; Welburn et al., 2010). These afore-mentioned studies coupled with the fact that Cdt1 closely resembles the Ndc80 and Ska complexes in being recruited to kinetochores where it binds to spindle MTs and contributes to kMT stabilization (Varma et al., 2012), prompted us to insinuate that Aurora B might be one of the essential kinases involved in regulating Cdt1-MT binding during mitosis.

A close examination of the Cdt1 sequence indeed revealed at least 7 Aurora B phosphorylation sites (T7, S73, S101, T102, S143, S265, T358) organized in a canonical Aurora B consensus motif [(R/K)_1-3_-X-(S/T)] (Meraldi et al., 2004) (**Fig. 3 A**). Five of these seven sites (S73, S101, T102, S143 and S265) have been reported as phosphorylation sites in the literature-curated database (www.phosphosite.org and www.phosida.org). Since, highly conserved S159 in Ska3 was clearly phosphorylated by Aurora B even though it did not match the canonical consensus (Chan et al., 2012), we also included 3 such sites (T270, T287 and T329) in Cdt1 that were conserved across the species but did not conform to the classical Aurora B sequence motif (**Fig. 3 A**).

**Figure 3.**
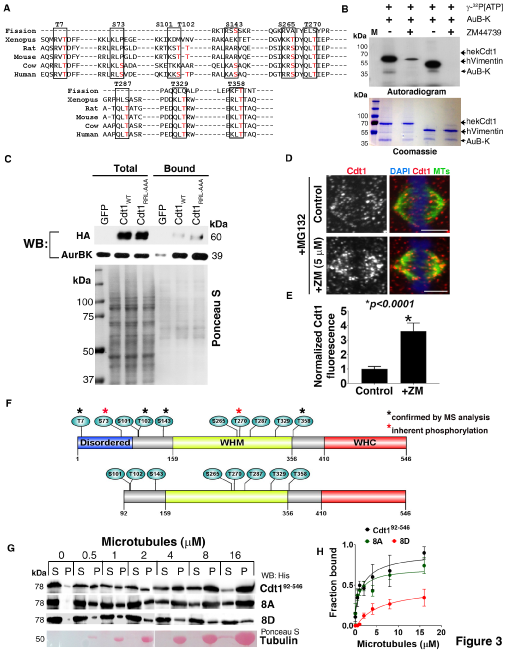
Aurora B Kinase targets Cdt1 for phosphorylation *in vitro* and affects Cdt1-MT binding. (A) Sequence alignment of Cdt1 from the indicated species using Clustal Omega. Highlighted in the red font are the potential Ser/Thr Aurora B sites. On the top, mentioned are the residues and their positions with respect to the full-length human Cdt1 (1-546 aa). (B) *In vitro* kinase assay with Aurora B alone or kinase plus ZM447439 inhibitor (10 μM) on HEK 293-purified Cdt1 or hVimentin (as a positive control). The autoradiogram (top) and Coomassie-stained gel (bottom) are shown. “M” denotes migration of molecular mass standards in kDa on a 12% SDS-PAG. (C) Pull down of Aurora B by HA/His-tagged Cdt1-WT or cy-Cdt1 mutant that blocks Cyclin A/Cdk binding [RRL (68-70) AAA], from thymidine synchronized and nocodazole arrested mitotic HeLa cell extracts. The pull down was performed using Ni+^2^-NTA agarose beads followed by immunoblotting with either anti-HA or anti-Aurora B (AurB-K) antibodies. 1% of the lysate was loaded as total protein. HEK-cells expressing GFP were used as a control. (D) HeLa cells were either treated with Aurora B inhibitor, ZM447439 (5 μM) for 1 h or left untreated (control) followed by release into MG132 (10 jM) containing medium in each case. The cells were immunostained with anti-Cdt1 (red), anti-tubulin (green) antibodies and counterstained with DAPI to mark the chromosomes (blue); Scale bar, 5 jm. (E) n=10 cells were quantified to obtain normalized Cdt1 fluorescence intensities on spindle MTs as depicted in D. (F) Schematic representation showing the full-length Cdt1 (546 aa) and its deletion variant devoid of N-terminal 92 aa generated in fusion with GFP tag for expression in bacteria. The following two proteins generated represent Aurora B phosphomimetic (8D, Ser/Thr substituted with Asp) and phospho-deficient (8A, Ser/Thr substituted with Ala) Cdt1 mutants. Black and red asterisks on the top, denote the confirmation of phosphosites by Mass spectrometry or residues that were inherently phosphorylated in Cdt1 obtained from HEK 293 cells, respectively. (G) Representative western blot is shown. Samples fractionated as Supernatant (S) and Pellet (P) analyzed by Western blot probed with antibody against 6×-His tag to detect Cdt1 and stained with Ponceau S for tubulin. (H) Quantification (Mean ± SD, n=3) showing MT co-sedimentation of purified Cdt1^92-546^ or the mutant proteins (1 μM each) with the indicated concentrations of taxol-stabilized MTs (in μM). The 95% CI values obtained for each fit were: *B_max_*=0.34- 0.65, *K_d_*=0.43-10.57 for 8D and *B_max_*=0.36-0.67, *K_d_*=0-3.22 for 8A.

To validate this hypothesis, we first sought to test if Cdt1 is a substrate for Aurora B kinase using an *in vitro* phosphorylation assay. Under the assay conditions, both myc-tagged full-length hekCdt1 (purified from HEK 293 cells and procured commercially) and Vimentin [an established Aurora B substrate (Goto et al., 2003)] were readily phosphorylated by Aurora B (**Fig. 3 B**). This phosphorylation was markedly reduced upon addition of ZM447439, an Aurora B inhibitor, attesting to its specificity. In tandem, hekCdt1 phosphorylated by Aurora B (using a non-radioactive ATP) was subjected to mass-spectroscopy for identification of phosphosites. The MS/MS spectra conclusively detected phosphorylation on four Aurora B consensus sites *viz* T7, T102, S143, and T358 uniquely/exclusively in the kinase treated hekCdt1; with 90% sequence coverage in two independent runs (**Fig. S2**). S73 and T270 were detected as phosphorylated even in the control hekCdt1 (without the Aurora B kinase); suggesting that these residues can potentially acquire phosphorylation *in vivo.* While the S101 and S265 localized on the peptides that were phosphorylated, the exact location of the modified residues could not be designated. The remaining two non-canonical sites T287 and T329 were unmodified.

Interestingly, Ni^+2^-NTA agarose-mediated affinity precipitation of HA/His-tagged Cdt1 (from thymidine synchronized and nocodazole-arrested mitotic cell extracts), but not GFP, led to the co-precipitation of Aurora B kinase as an interacting partner (**Fig. 3 C**). Moreover, the kinase was also co-precipitated with the HA-tagged Cdt1 mutant [RRL(68-70)AAA], which was incompetent to bind Cyclin/Cdk, equally efficiently as with the wild-type Cdt1; implying that the interaction between Cdt1 and Aurora B in mitosis is independent of Cdk-phosphorylation (**Fig. 3 C**).

To test the significance of Aurora B phosphorylation on the MT-binding of Cdt1 *in vivo,* we treated HeLa cells with Aurora B inhibitor (ZM 447439) and assessed the localization of Cdt1 on mitotic spindle MTs. We observed a 3.6-fold increase in spindle MT-localized Cdt1 in ZM-treated cells compared to the untreated control cells (**Fig. 3, D and E**). Consistent with this finding, upon mutating the 8 putative Aurora B Ser/Thr residues (8D) to Asp in Cdt1^92-546^ in an attempt to mimic constitutively phosphorylated Cdt1 (**Fig. 3 F**), we observed ~2.5-fold reduction in the apparent MT binding as evidenced by an increase in the *K_d_* from 2.03 ± 0.99 μM for the Cdt1^92-546^ to 5.50 ± 2.41 μM for the 8D mutant (**Fig. 3, G and H**). Substitution of the residues to non-phosphorylatable Ala (8A) did not impact the MT binding with *K_d_* value comparable to the parental Cdt1^92-546^.

### Aurora B-phosphomimetic mutations in Cdt1 affects kMT stability and mitotic progression

To assess the role of Aurora B-mediated phosphorylation for Cdt1-MT binding *in vivo,* we investigated the function of the Aurora B phosphomimetic and non-phosphorylatable Cdt1 mutants in HeLa cells using an experimental strategy that had been established previously. This approach involves siRNA-based knockdown and rescue of Cdt1 function in double thymidine synchronized cells specifically in the G2/M phase of the cell cycle to ensure that Cdt1 function in replication licensing during G1 phase is unperturbed (Varma et al., 2012). We first generated HeLa cells stably expressing HA-tagged wild-type and phosphomutant Cdt1 using a retroviral transduction-based approach. The stability of kMTs in cells which were depleted of endogenous Cdt1 but rescued with Cdt1 wild-type or phosphorylation site mutants was tested. Upon cold treatment, the expression of Cdt1-WT rescued kMT stability as opposed to that of the phosphomimetic mutant (Cdt1-10D) or the vector control (simulates Cdt1-depleted state) (**Fig. 4, A and B**). However, the non-phosphorylatable mutant (Cdt1-10A) not only maintained stable kMT attachments but the quantitation of fluorescence intensities of spindle MTs suggested that the kMTs were hyper-stable in this case (**Fig. 4, A and B**). Under our conditions, efficient knockdown of endogenous Cdt1 was achieved while the expressed siRNA resistant HA-tagged Cdt1 proteins were retained (**Fig. 4 C top and bottom panels**). The cells expressing either the WT or mutant proteins after endogenous Cdt1 depletion had apparently normal spindle structures at 37°C (**Fig. S3 A**) and no defect in localization of Hec1 to kinetochores (**Fig. S3 B**).

**Figure 4.**
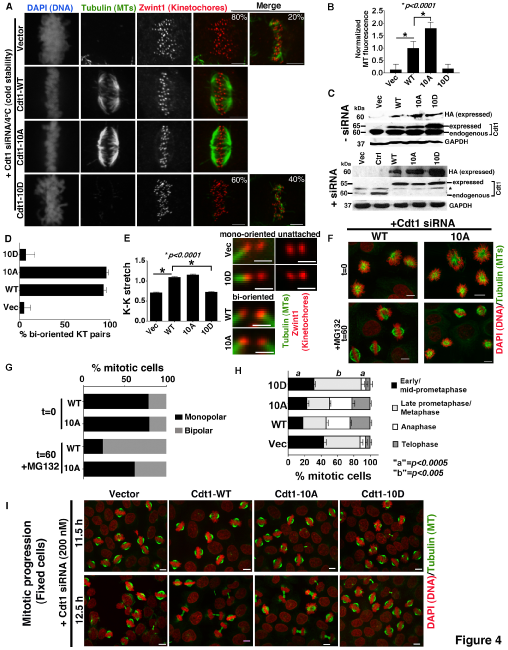
Aurora B phosphorylation of Cdt1 regulates kinetochore microtubule stability and mitotic progression. (A) Synchronized HeLa cells treated with siRNA against endogenous Cdt1 in G2/M phase and rescued with stably expressing siRNA-resistant Cdt1-WT, 10D (phosphomimetic), 10A (unphosphorylated) proteins or Vector control (abbreviated as Vec, simulating Cdt1 depleted state); were stained with antibodies against Zwint1 in red (1:400 dilution, kinetochore marker), MTs in green (1:500 dilution, spindle marker) and DAPI in b/w marking the chromosomes. Cells were placed at 4°C for 15 min before being fixed and stained. Scale bar, 5 μm. (B) A bar graph showing the quantification of MT staining intensity in each case after background subtraction (n=10 cells). (C) HeLa cells treated with siRNA against endogenous Cdt1 rescued with stably expressing siRNA-resistant Cdt1-WT, 10D (phosphomimetic), 10A (phosphodefective) proteins (with C-terminal HASH tags) or Vector control (Vec) were lysed using 2×-laemelli buffer and electrophoresed on 10% SDS-PAG. Representative Western blots from three independent experiments of synchronized HeLa cell lysates probed for expression levels of the indicated proteins with (bottom) or without (top) Cdt1 siRNA (200 nM). “Expressed” and “Endogenous” represent ectopically expressed (with HASH tags) and endogenous (without tag) Cdt1, respectively. Asterisk (*) in the bottom panel denotes detection of a non-specific band reactive with the Cdt1 antibody. The same blot was re-probed with GAPDH (1:8000) as a sample recovery control. (D) Quantification of the % of bi-attached, bi-oriented kinetochores in each of the conditions as indicated. Kinetochore marker, Zwint1, is in red and MTs is in green. (E) Inter-kinetochore (K-K) stretch was calculated between the two kinetochore pairs in each case and is plotted (*n*=115 pairs; 10 cells). Cropped images of representative kinetochore pairs are also provided to indicate the status of their kMT attachment and K-K stretch; scale bars, 1 jm. (F) STLC washout assay was carried out in double thymidine synchronized HeLa cells, treated with siRNA against endogenous Cdt1 and rescued with either Cdt1-WT or Cdt1-10A followed by treatment with STLC for 2 h. The cells were washed with and released into medium containing MG132 and were fixed either immediately after the initiation of the washout (t=0) or after 1 h (t=60), followed by fixation and staining. DAPI is pseudo-colored in red to mark chromosomes and Tubulin in green. (G) A bar graph showing the quantification of monopolar (early prometaphase) *vs.* bipolar spindle (metaphase) structures at the indicated time points after STLC washout. Cells from two coverslips in two independent experiments were counted; n=2114 and 2659 for WT and 10A, respectively for t=60 and n=565 and 785 for WT and 10A, respectively for t=0. The data is plotted as % mitotic cells with monopolar or bipolar spindle structures on the Y-axis. (H) Mitotic progression (metaphase to anaphase transition) was assessed in HeLa cells stably expressing vector, Cdt1-WT, Aurora B Cdt1-phosphomimetic (10D) or phospho-defective (10A) mutants at 5 and 12.5 h after release from thymidine treatment in the presence of Cdt1 siRNA. The number of cells in each stage of mitosis were counted and plotted from three different coverslips of two independent experiments to illustrate the nature of mitotic delay/arrest in Cdt1-10D mutant-expressing cells as compared to cells rescued with Cdt1-WT or Cdt1-10A expression (n=600 cells; “a” and “b” represent the statistical significance obtained using two-sided unpaired non-parametric students *t* test performed on the WT *vs* 10D in the indicated stages of mitosis). (I) Representative images are shown from H; DAPI pseudo-colored in red marks the chromosomes and antibody against tubulin stained the MTs green.

Approximately 60% of the cells analyzed in Cdt1-10D and 80% in vector control had no discernible MT structures after cold exposure; in the remaining cells that showed extant MT traces, the MTs were not attached to the kinetochores (**Fig. 4 A**, last panels in each vector and 10D). Quantification of the bi-oriented kinetochores that made contact with cold-stable MTs from opposite spindle poles revealed ~90% decrease in cells expressing phosphomimetic Cdt1 as compared to cells expressing wild-type or phospho-defective Cdt1 (**Fig. 4 D**). The inter-kinetochore distance (K-K stretch), which serves as another indicator of the strength of kMT attachment, was also measured in each case as another parameter. The average K-K stretch in cells expressing Cdt1-WT and Cdt1-10A was 1.11 ± 0.09 μm and 1.16 ± 0.15 μm, respectively (**Fig. 4 E**); implying kMTs from opposite spindle poles forming robust contact with sister kinetochores. In contrast, the Cdt1-10D mutant exhibited a significant reduction in the distance between sister kinetochores (0.73 ± 0.10 μm) similar to the vector control, 0.72 ± 0.08 μm indicating weakened load-bearing kMT attachments between kinetochore pairs (**Fig. 4 E**). Magnified images of kinetochore pairs are also shown to underscore the observed effects (reduction of the K-K distance and occurrence of mono-oriented or unattached KT pairs) in Aurora B phosphomimetic mutant and the vector control (**Fig. 4 E**).

In order to conclusively address if expression of Cdt1-10A led to the production of hyper-stable kMT attachments, we carried out STLC (analogous to Monastrol, an inhibitor of Kinesin Eg5) washout assay to test the ability of cells to correct syntelic attachments (Kapoor et al., 2000). The data revealed that 1 h post-drug washout, only ~38% of cells expressing non-phosphorylatable Cdt1 mutant (10A) had attained bipolar spindle structures; in contrast to ~76% of cells rescued with the WT-Cdt1 protein (**Fig. 4, F and G**). Based on the substantial delay in bipolarization of the mitotic spindles and accumulation of either monopolar spindles or early prometaphase like spindle structures in 10A rescued cells, it is evident that there is some degree of kMT hyper-stability associated with the expression of the Cdt1-10A mutant as compared to cells rescued with WT-Cdt1.

Progression through mitosis was monitored in each case by fixed-cell analyses. Cells were fixed at two different time points after release from thymidine arrest; mitotic onset (11.5 h) and anaphase onset (12.5 h) (Varma et al., 2012). The mitotic progression of cells expressing Cdt1-10D was severely disrupted. Similar to that observed for vector control after the depletion of endogenous Cdt1 (Varma et al., 2012), the Cdt1-10D expressing cells exhibited a phenotype resembling a mid to late pro-metaphase arrest at the latter time point (**Fig. 4, H and I**). In this case, majority of the cells had chromosomes that were partially aligned around a vaguely defined metaphase plate (**Fig. S3 A**), while a smaller fraction of cells exhibited substantially higher levels of chromosome misalignment (data not shown). These phenotypes were strikingly different from the cells expressing the Cdt1-10A mutant, which underwent anaphase onset and cytokinesis similar to the cells rescued with Cdt1-WT.

Although we did not demonstrate that these 10 sites in Cdt1 could be phosphorylated by Aurora B kinase *in vivo*, mass spectrometric data unequivocally confirmed at least 4 target residues (T7, T102, S143, and T358) that were phosphorylated by Aurora B *in vitro.* Therefore, in order to evaluate whether mutating the 4 identified sites is sufficient enough to recapitulate some or all of the observed phenotypes (especially perturbation of kMT stability and mitotic progression), we generated HeLa cells stably expressing siRNA resistant versions of HA-tagged phosphomimetic (4D) and non-phosphorylatable (4A) mutant Cdt1 proteins. Employing the above-mentioned siRNA-based-knockdown-rescue approach, analysis of kMT cold stability and fixed-cell mitotic progression analyses were conducted for these mutants as well. Results from the cold stability experiment indicate that cells rescued with the expression of Cdt1-4D mutant after the depletion of endogenous Cdt1 still exhibited substantial loss of kMT stability as compared to the cells rescued with WT-Cdt1 even though the effects were not as severe as those observed in cells expressing Cdt1-10D mutant (**Fig. 5 A**). Consequently, the incidence of unattached or mono-oriented kinetochores after Cdt1-4D expression was intermediate/partial with respect to Cdt1-WT and Cdt1-10D expressing cells. Interestingly, the cells expressing Cdt1-4A also displayed significant increase in the stability of kMTs when compared to the Cdt1-WT, but to a significantly lesser extent than the cells expressing the Cdt1-10A mutant (**Fig. 5 B**). Mitotic progression analysis corroborates the above data as cells rescued with Cdt1-4A/4D exhibited phenotypes of intermediate severity between the Cdt1-WT expression and the more penetrant, 10A/10D mutant expression (**Fig. 5, C and D**). As previously described, the cells were fixed at two different time points; mitotic onset (11.5 h) and anaphase onset (12.5 h). While fewer mitotic cells expressing Cdt1-4A were able to enter anaphase as compared to those expressing Cdt1-10A at 12.5 h; there was a significant increase in the number of mitotic cells expressing Cdt1-4D that were able to enter anaphase as compared to the Cdt1-10D expressing cells (**Fig. 5, C and D**). In support of the *in vivo* analysis, microtubule co-pelleting assay was also performed with the 3D mutant where the three out of four identified sites in Cdt1^92-546^ were substituted with Asp to mimic constitutively phosphorylated Cdt1 (**Fig. 5 E**). The apparent *Kd* of the 3D mutant for MTs was calculated to be 3.27 ± 0.66 as opposed to 1.85 ± 0.34 μM for the Cdt1^92-546^; indicating ~1.5-fold reduction in the MT binding ability of the 3D mutant (**Fig. 5, F and G**).

**Figure 5:**
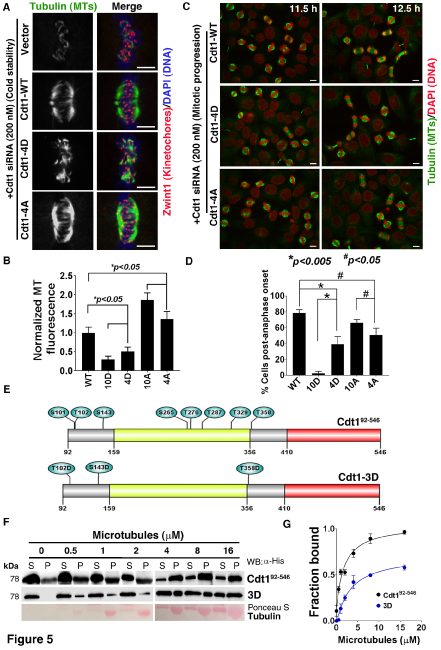
Aurora B phospho-mutants of Cdt1 at the four sites identified by mass spectroscopic analysis partially affects kinetochore microtubule stability and mitotic progression. Synchronized HeLa cells treated with siRNA against endogenous Cdt1 in G2/M phase and rescued with stably expressing siRNA-resistant Cdt1-WT, 4D (phosphomimetic), 4A (unphosphorylated) proteins or Vector control (simulating Cdt1 depleted state) were subjected to 4°C treatment for 15 min before being fixed and stained. (A) Representative images of cells from C, immunostained with antibodies against Zwint1 (kinetochore marker) in red, tubulin (MTs) in green or in b/w on the left of the merged images, and DAPI in blue marking the chromosomes. Scale bar, 5 μm. (B) A bar graph showing the quantification of spindle MT staining intensity after cold treatment in each case after background subtraction (n=7). (C) Mitotic progression (metaphase to anaphase transition) was gauged in HeLa cells stably expressing Cdt1-WT, Cdt1-10D, Cdt1-10A, Cdt1-4D, and Cdt1-4A mutants at 11.5 and 12.5 h after release from thymidine treatment in the presence of Cdt1 siRNA (200 nM). Representative images are shown from E; DAPI pseudo-colored in red marks the chromosomes and antibody against tubulin stained the MTs green; Scale bar, 10 μm. (D) The number of cells in each stage of mitosis were counted and plotted from three different coverslips to illustrate the difference in mitotic delay in Cdt1-4A/4D mutant-expressing cells as compared to cells rescued with Cdt1-WT or Cdt1-10A/10D expression (n=300); * and # represent the statistical significance amongst the indicated groups obtained using two-sided unpaired non-parametric students *t* test. (E) Schematic representation showing the 3 indicated Aurora B sites mutated to generate a 3D phosphomimetic mutant in the Cdt1^92-546^ parental background. (F) Representative western blot from a MT-cosedimentation experiment performed with Cdt1^92-546^ and Cdt1-3D mutant is also shown. Samples fractionated as Supernatant (S) and Pellet (P) analyzed by Western blot probed with antibody against 6×-His tag to detect Cdt1 (WT or 3D mutant) and stained with Ponceau S for tubulin. (G) Quantification (Mean ± SD, n=3) showing MT co-sedimentation of purified Cdt1^92-546^ or the mutant proteins (1 μM each) with the indicated concentrations of taxol-stabilized MTs (in μM). The 95% CI values obtained for each fit were: *B_max_*=0.8-1.0, *K_d_*=1.22-2.90 for Cdt1^92-546^ and *B_max_*=0.67-0.85, *K_d_*=2.25-4.9 for 3D mutant.

We carried out live cell imaging to substantiate our data from fixed-cell mitotic progression s. However for this analysis, we chose Cdt1-10A and −10D mutants since they yielded a more complete and potent phenotype. The vector control and cells rescued with Cdt1-WT or −10A or −10D Aurora B mutant versions were subjected to live imaging after the knockdown of endogenous Cdt1 (**Fig. 6 A and Movies S4-S7**). Hoechst was used to mark the chromosomes prior to the initiation of live imaging. Since there was no noticeable delay in the alignment of the majority of chromosomes in any of the 4 samples, we quantified the time elapsed between the initial establishment of the metaphase plate and the final fate of each of the imaged mitotic cell. We find that ~100% of the Cdt1-WT rescued mitotic cells underwent normal anaphase onset without any delays (n=26/27, **Movie S4**) while >80% of the vector control (n=22/27, **Movie S5**) and the Cdt1-10D rescued cells (n=23/28, **Movie S6**) exhibited a prolonged mitotic arrest with a metaphase-like chromosome alignment. In the latter two conditions, the aligned state of the chromosomes was a) maintained for an extended period of time until the last time point of live imaging (outcome 1), or b) followed by anaphase onset with chromosome missegregation events (outcome 2), or c) disrupted by chromosomal de-condensation and reversion to an interphase-like stage (outcome 3). More importantly, vector control and the Cdt1-10D expressing mitotic cells, on an average took ~3 times longer (~200 min) to reach their final fate (outcomes 1, 2, and, 3 above) with ~15% cells never progressing into anaphase (outcome 1) (**Fig. 6 B and Movies S5 and S6**). Even though 10A mutants progressed through mitosis comparable to the WT-rescued cells, >50% of these cells (n=15/28, **Movie S7**) underwent chromosome missegregation events during anaphase, which we speculate is a consequence of hyperstable kMT attachments.

**Figure 6:**
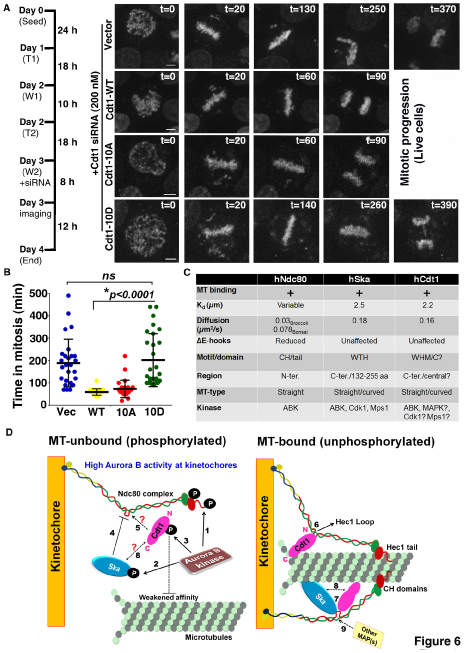
Live cell imaging analysis of cells rescued with Aurora B phosphomimetic Cdt1 mutant reveals a severe delay in mitotic progression. (A) Scheme of the time course and still images obtained at the indicated time points during the imaging of live mitotic progression in different stable cell lines rescued with either the plasmid vector (analogous to Cdt1 depletion) or by WT or Aurora B mutant Cdt1 expression. Chromosomes are marked with Hoechst dye. (Also, see Supplementary movies S4-S7). (B) Quantification of (A). The time (in min) elapsed between the establishment of a metaphase plate and the final fate of each of the imaged mitotic cell, which included delayed anaphase onset or a prolonged mitotic arrest, was plotted on the Y-axis for each of the sample imaged live as indicated. (C) Comparative tabulation of MT-binding parameters and phospho-regulation of relevant kinetochore associated proteins from vertebrates. (D) A working model depicting the role of Cdt1 in providing an additional kMT-attachment site besides the CH and the N-terminal tail domain of the Hec1 subunit of the Ndc80 complex. The calponin homology (CH) domains in Hec1 and Nuf2 along with the Hec1 unstructured tail bind MTs. The unstructured ~40 aa loop region of Hec1 recruits Cdt1 by interacting with the N-terminal 1-320 aa of Cdt1 (Varma et al., 2012). This induces a conformational shift in the Ndc80 complex in a manner that is dependent on Cdt1 binding to MTs. In prometaphase when kinetochores are not bi-oriented, Aurora B kinase is active to destabilize the erroneous attachments by phosphorylating several kinetochore proteins including Ndc80 (1), Ska (2), and Cdt1 (3) shown in the present study, all of which bind to MTs. Upon phosphorylation, these proteins exhibit reduced affinity for MTs. As observed for the Ska complex which cannot interact with the KMN network effectively upon its phosphorylation by Aurora B (4); whether Cdt1~P can dock on to the Hec1 loop (5) with similar efficiency as unphosphorylated Cdt1 is still undetermined. In metaphase, when the Aurora B gradient diminishes at kinetochores, Cdt1 becomes proficient to binds to MTs (6), thus providing an additional site for kMT attachment besides the CH and the tail domains of Ndc80. Whether Cdt1 can interact with the Ska complex in its free or loop-bound state and if this interaction is dependent on their phosphorylation status (7 and 8) is yet to be determined. The model also depicts the possibility of interaction of other MAPs with the Hec1 loop either alone or with the help of Cdt1 or Ska or both (9), as shown using dashed arrows.

## Discussion

In this study, we demonstrate that the DNA replication licensing protein Cdt1, which is recruited to mitotic kinetochores by the Ndc80 complex (Varma et al., 2012), also binds to MTs directly in a manner that its affinity is controlled by Aurora B kinase phosphorylation. The affinity of Cdt1 for MTs was found to be comparable with other relevant kinetochore MAPs such as chTOG (Spittle et al., 2000), Ska (Schmidt et al., 2012), the KMN network and the Ndc80 complex (Cheeseman et al., 2006; Ciferri et al., 2008) computed using similar approaches. Lack of contribution from the Cdt1’s N-terminal 92 aa towards MT-binding indicates that the mechanism by which Cdt1 binds to MTs is distinct from the Hec1 subunit of Ndc80, where a positively charged N-terminal 80 aa long unstructured tail is essential for proper MT-binding of the Ndc80 complex (Guimaraes et al., 2008). The fact that the C-terminus of Cdt1 (410-546 aa) was not able to bind to MTs alone as effectively as in the presence of N-terminus is reminiscent of the pattern obtained with the Ska complex; wherein the removal of C-termini from either Ska1 or Ska3 impaired MT-binding but at the same time, the purified C-terminal domains of Ska1 or Ska3 by itself did not associate with MTs (Jeyaprakash et al., 2012). This suggests that the N-terminus of Cdt1 (1-410 aa) may be inert for MT binding by itself but might facilitate the folding or positioning of the C-terminus of Cdt1 to allow MT binding. Moreover, we predict that the C-terminal (WHC) or centrally located (WHM) domains may be required in combination for efficient binding of Cdt1 to MTs. These observations also suggest that the WHD domain could serve as one of the conserved MT-binding domains in MT-binding proteins as has already been demonstrated for the Ska complex (Schmidt et al., 2012; Abad et al., 2014). Further, it is interesting that the N-terminal region of Cdt1 although could not bind to MTs, it was competent enough to interact with Hec1 and thus it seems that Cdt1 modular functions (Hec1- and MT-binding) are independent of each other even though Ndc80 loop is essential for recruiting Cdt1 to the kinetochores.

Additionally, our study demonstrates for the first time that Cdt1 is a substrate for Aurora B kinase and provides evidence for a physical interaction between the two proteins *in vivo.* However, in the pull down experiment, we found that the amount of tagged Cdt1 retrieved/bound (**Fig. 3 C**, IB: HA, Lanes 5 and 6) was much less when compared to the “total” protein expressed (Lanes 2 and 3). This is because the polyhistidine tag used to retrieve Cdt1, binds to nickel resin very specifically but extremely poorly in the incompatible buffer conditions used, essentially to preserve and retain Cdt1 binding partners. Nonetheless, endogenous Aurora B kinase was associated significantly above the background control in each case. In addition to Aurora B kinase, Cdt1 has been shown to be phosphorylated by Cyclin-dependent kinase 1 (Cdk1) during the S-phase at Threonine 29 (Fujita, 2006) and by Mitogen-activated protein (MAP) kinases p38 and c-Jun N-terminal kinase (JNK) during G2/M phase (Chandrasekaran et al., 2011). Fairly recently, Ska3 has also been shown to undergo Cdk1-mediated phosphorylation that influences its interaction with the Ndc80 loop domain (Zhang et al., 2017). Mps1 kinase was shown to phosphorylate both Ska3 (Maciejowski et al., 2017) and the yeast Ndc80 complex (Kemmler et al., 2009); in addition to a physical interaction with human Ndc80 (Nilsson, 2015). For Hec1, although Aurora B is regarded as the master regulator, the involvement of Aurora A kinase in late mitosis has lately been uncovered (DeLuca et al., 2018). With these ever-growing evidences of spatio-temporal integration of several mitotic kinases to fine-tune kMT stability, attachment error correction and mitotic progression; we envision a similar interplay of kinases in regulating mitotic functions of Cdt1, besides Aurora B.

Nevertheless, we extended our study to understand the implications of Aurora B kinase mediated phosphorylation on MT-binding ability of Cdt1. Indeed, our data demonstrated a correlation between the phosphorylation of Cdt1 at the 10 potential Aurora B sites and MT binding. While the phosphomimetic Cdt1-10D mutant was severely compromised to sustain stable kMT attachments, the Cdt1-10A mutant on the contrary exhibited enhanced attachment stability. However, this hyperstability did not translate into any discernible chromosome alignment defects unlike those observed after Hec1-9A mutant expression (DeLuca et al., 2011). This could be attributed to the specific role of Cdt1 during later stages of mitosis, i.e. at metaphase where it facilitates stabilization of kMT attachments. This is in contrast to the Hec1 where Aurora B-mediated Hec1 phosphorylation is actively involved in error correction during early mitosis. Thus, it is reasonable that hyperstable kMT attachments would lead to defective chromosome alignment in Hec1-9A mutant but in the case of Cdt1, it only translates to erroneous chromosome segregation later during anaphase onset.

In an attempt to identify the exact functional Aurora B site(s) that can sufficiently mimic the observed phenotypes, we focused on only the four residues that were unambiguously identified by mass-spectroscopic analysis. While our results confirm the vital contribution of those 4 Aurora B sites; phosphorylation on some or all of the other remaining sites seems requisite in order to attain the complete gamut of the phenotype pertinent to kMT stability. In light of the severity of phenotype obtained with the 10D mutation, the possibility of a complete loss of mitotic function for this protein cannot be completely ruled out. It is intriguing that these 10 predicted Aurora B sites are located within the N-terminal unstructured and central WHM (1-410 aa) of Cdt1 that did not show direct association with MTs but somehow seem to contribute to efficient Cdt1-MT interaction (**Fig. 1, G and I**). Since many of these sites are clustered in regions which are predicted to be disordered, we surmise that Aurora B phosphorylation might induce a conformational change in Cdt1 to regulate its MT-binding ability. This mode of regulation can exist in addition to or in parallel to the manner in which Aurora B phosphorylation generally modulates the affinity of binding; i.e. by interfering with the electrostatic interactions through introduction of multiple negative charges. Unlike Hec1, where multiple Aurora B target sites are grouped in tandem within the small unstructured N-terminal tail, sites within Cdt1 are distributed throughout. Thus we propose that one or both of the following could have elicited destabilization of kMT attachments leading to delayed mitotic progression upon Aurora B phosphorylation of Cdt1; (a) addition of multiple negative charges and/or (b) a conformational shift.

The behavioral parameters and regulatory aspects of the key MT-binding proteins which participate in stabilizing kMT attachments (including Cdt1) at the vertebrate kinetochores are summarized in **Fig. 6 C**. Based on our observations, we propose a model where kinetochore-localized Cdt1 provides another MT-binding site for the Ndc80 complex at its loop domain, thus shedding light on the importance of Cdt1 and the loop in kMT attachments (**Fig. 6 D**). We envisage that the loop domain of the Ndc80 complex either alone or in conjunction with bound Cdt1 might also coordinate with other kinetochore MAPs to confer stronger attachments. Thus, future studies will be required to delineate the details of hierarchical recruitment of kinetochore components by the Ndc80 loop. Our work, while providing an essential missing link in the series of events that take place during the stabilization of kMT attachments in mitosis (**Fig. 6 D**), also opens up a number of stimulating questions such as: (a) whether the Ndc80 complex has any synergistic influence on Cdt1 binding to MTs; (b) does Ska and Cdt1 coordinate for robust kMT attachments and if so, do these proteins dock on to the loop as a pre-formed complex or do they interact after they are independently recruited at the loop; (c) can Ska, Ndc80 and Cdt1 generate a ternary complex for efficient MT binding; (d) does chTOG, another Ndc80-recruited kinetochore MAP influence localization of Cdt1 and its interaction with MTs; (e) is there a cross-talk and/or interplay of several other kinases besides Aurora B kinase to fine-tune the phosphoregulation of Cdt1 in mitosis; (f) how phosphorylation of Cdt1 with one or many mitotic kinases influences its localization to kinetochores and whether it affects recruitment of other kinetochore-associated proteins directly or indirectly? Answers to these questions will have major impact on the mitosis field in the quest for novel mechanisms that link mitotic kinetochores to spindle MTs and drive accurate chromosome segregation.

## Acknowledgments

We thank Dr. Jeanette Cook for the HA-tagged Cdt1 expression plasmid and for valuable intellectual input; Dr. Sarah Rice for constructive suggestions and discussions; Drs. Laimonis A Laimins and Kavi Mehta for providing help with the radioactive experiments; Drs. Arshad Desai and Dhanya Cheerambathur for providing Hec1::Nuf2-His tagged dimeric complex, Northwestern Genomics Core for DNA sequencing services; Suchithra Seshadrinathan and Dr. Anita Varma for technical support. This work was supported by NCI grant R00CA178188 to DV; NINDS grant R00NS089428 and NIGMS grant R35GM124889 to RJM.

The authors declare no competing financial interests.

## Author contributions

S.A. and D.V. designed and performed majority of the experiments. K.P.S. assisted in Cdt1 purification, carried out structural and computational studies and provided intellectual input. Y.Z. performed the Cdt1-Aurora B kinase pull down experiment. A.S. performed endogenous Cdt1 antibody staining. R.J.M. performed single molecule TIR-FM experiments. S.A., R.J.M. and D.V. analyzed the data. S.A. and D.V. wrote the manuscript with input from all the authors.

## Abbreviations

kMT: kinetochore-microtubule
MT(s): microtubule(s)
Knl1: Kinetochore null 1
Ndc80: Nuclear division cycle 80
Mis12: Missegregation 12
CH: Calponin homology
MAP(s): Microtubule associated protein(s)
Ska: Spindle and kinetochore-associated protein
TOG: Tumor overexpressed gene
MTBD: microtubule binding domain
WH: Winged helix
WH(M/C): Winged helix middle or C-terminal

## Materials and methods

### Chemicals, reagents, cell lines and bacterial strains

Bacterial strains for cloning and expression were grown in Luria-Bertani (LB) broth or agar supplemented with 100 μg/ml ampicillin or 50 μg/ml kanamycin as appropriate. Enzymes for cloning were purchased from New England Biolabs (Beverly, MA). Custom oligonucleotides and G-blocks were from Integrated DNA Technologies (Coralville, IA). Sanger DNA sequencing was conducted at the Northwestern University Genomics core facility.

### DNA manipulations: cloning and mutagenesis

Cdt1 gene sequence was codon optimized and was synthesized as a linear DNA strand (G-block, IDT) for subsequent cloning procedures. The largest Cdt1 fragment (92-546 aa), deletion variants (92-232 aa, 92-410 aa, 410-546 aa) and Aurora B phosphomimetic mutants (Ser101Asp, Thr102Asp, Ser143Asp, Ser265Asp; Thr270Asp, Thr287Asp, Thr329Asp, Thr358Asp abbreviated as 8D and Ser101Asp, Ser143Asp, Thr358Asp abbreviated as 3D) and non-phosphorylatable mutants (Ser101Ala, Thr102Ala, Ser143Ala, Ser265Ala; Thr270Ala, Thr287Ala, Thr329Ala, Thr358Ala abbreviated as 8A); were cloned in fusion with the N-terminal GFP and 6×-His tags using SspI and Gibson cloning in a commercial plasmid procured from Adgene (pET His6 GFP TEV LIC cloning vector (1GFP), plasmid #29663).

For stable transfections, the genes encoding Cdt1 with mutations that represent Aurora B phosphomimetic (Thr7Asp, Ser73Asp, Ser101Asp, Thr102Asp, Ser143Asp, Ser265Asp; Thr270Asp, Thr287Asp, Thr329Asp, Thr358Asp) and non-phosphorylatable (Thr7Ala, Ser73Ala, Ser101Ala, Thr102Ala, Ser143Ala, Ser265Ala; Thr270Ala, Thr287Ala, Thr329Ala, Thr358Ala) versions, abbreviated as 10D and 10A, respectively harboring C-terminal 2×-HA, Strep-tag, 12×-His tag were cloned in pQCXIP plasmid using custom gene synthesis services (Thermofisher Scientific). In addition, Aurora B phosphomimetic (Thr7Asp, Ser101Asp, Ser143Asp, Thr358Asp) and non-phosphorylatable (Thr7Ala, Ser101Ala, Ser143Ala, Thr358Ala) versions, abbreviated as 4D and 4A; with the same tags as mentioned above were also custom synthesized by Gene Universal Inc.

### *In silico* analyses

#### Secondary Structure Prediction

The protocol was based on the work of Seeger *et al.* (Seeger et al., 2012). The full sequence of human Cdt1 (Uniprot ID: Q9H211) was input into ten different secondary structure prediction servers to reduce the bias from any one program. The prediction of disordered regions is based on amino acid composition, charge/hydropathy index, known disordered sequences from crystal/NMR structures, and other empirical methods. The servers included were PSIPRED, IUPRED, PONDR, FoldIndex, s2D, ESpritz, PROFsec, MFD2p, DISCoP, and YASPIN. Relatively stringent cutoff of >75% within each algorithm was used to classify a residue as “disordered.” The results of each server were converted to a binary output of “folded/unknown” or “disordered” (0 or 1 respectively). The scores were summed and plotted as a function of amino acid number.

#### Homology Modeling

The full sequence of human Cdt1 (Uniprot ID: Q9H211) was input into the Phyre2 (Kelley et al., 2015), I-TASSER (Roy et al., 2010), and SWISS-MODEL (Schwede et al., 2003) homology modeling servers. Models were aligned using the “cealign” plugin on Pymol. The C-terminal winged helix domain was chosen to align all models. Crystal structures from human WHM domain (PDB ID:2WVR) (De Marco et al., 2009) and mouse WHC domain (PDB: 2KLO) (Khayrutdinov et al., 2009) were used as templates for the winged helix domains. Pymol was used to generate protein structure images and the final figures were generated in Adobe Illustrator.

#### Recombinant Protein Expression and Purification

For MT co-pelleting assays, the fusion proteins along with the GFP protein (as a control) were purified from the soluble fraction after over-expression in BL21(λDE3) at 23 °C for 18 h in LB medium, using nickel-affinity chromatography. The proteins purified to 85% purity were dialyzed in 20 mM Tris-HCl, pH 8.0 with 300 mM NaCl and stored in −20 °C until further use. Full length Cdt1 (1-546 aa) was purified as a GST tagged protein using pGEX-5X-1 followed by tag removal as per manufacturer’s instructions.

Native tubulin was purified from porcine brains as previously described (Williams and Lee, 1982), flash frozen, stored in aliquots in liquid nitrogen, and clarified of aggregates by centrifugation at 100,000 × *g* prior to use.

Full-length human Cdt1 (1-546 aa) purified from HEK293 cells with a C-terminal Myc/DDK tag dialyzed in 25 mM Tris-HCl, pH 7.3, 100 mM glycine and 10% glycerol was purchased from Origene (TP301657).

For TIRF assays, GFP tagged Cdt1, abbreviated as Cdt1^92-546^ was induced in BL21(λDE3) cells at 30 °C for 5 h in enriched terrific broth. Cells pellets were flash frozen in liquid nitrogen and stored at −80 °C. Cell pastes were lysed via sonication, and spun at 35,000 rpm for 45 min. The soluble fraction was incubated with 6 mL of 50% cobalt IMAC resin (TALON, Clontech) for 1 h at 4 °C. The beads were washed with 100 mL of wash buffer (25 mM HEPES pH 7.4, 500 mM NaCl, 2.5 mM imidazole, 5% sucrose w/v and 0.5 mM TCEP) and eluted with 15 mL of elution buffer (wash buffer + 500 mM imidazole). The eluted protein was concentrated to approximately 2.5 mL and injected over a Superdex 200 16/60 SEC column (GE Healthcare) in storage buffer (25mM HEPES pH 7.4, 500 mM NaCl, 0.5 mM TCEP, 5% sucrose w/v). Purified, folded Cdt1 eluted at approximately 65 mL and was monodisperse. The final protein was ~95% pure via SDS-PAGE. Protein was flash frozen in liquid nitrogen and stored at −80 °C until use.

#### Microtubule co-pelleting assay

For microtubule-binding assay, tubulin was polymerized and stabilized in general tubulin (GT) buffer, BRB80 (80 mM PIPES pH 6.8, 1 mM MgCl_2_, 1 mM EGTA) with 100 mM GTP, 20 μM Paclitaxel/taxol and 66 % glycerol (assay buffer). The proteins (diluted in BRB80 buffer containing 1 mg/ml BSA) were pre-cleared of any aggregates or debris by ultracentrifugation at 70,000 rpm at room temperature for 30 min. The taxol-stabilized MTs (in varying micromolar concentrations) were incubated with indicated concentration of pre-cleared protein(s) or 2 μM of GFP/GST alone to a final reaction volume of 40 μl. Reactions were incubated at room temperature for 20 min and ultra-centrifuged for 30 min at 60,000 rpm in a Beckman TLA100 rotor at 25 °C. The binding was also carried out in the presence of salt by adding 300 mM NaCl to the assay buffer. Pellets (P) and supernatants (S) were separated carefully, equal volumes of SDS sample buffer was added and then samples were run on SDS-PAG and analyzed by Western blotting with anti-His Monoclonal antibody (1:5000, Abcam) or Cdt1 polyclonal IgG raised in rabbit (1:2000, H-300, sc-28262, Santa Cruz Biotech). Immunoblots were scanned onto Adobe Photoshop and any manipulations to the brightness were applied to the entire image. Densitometry quantification of the bands representing the arbitrary amount of protein in supernatants and pellets was done using ImageJ software (National Institutes of Health, Bethesda, MD). The percentage binding was measured by calculating the ratio of protein in the pellet (P) and the total protein (S+P). The fraction of protein bound was then plotted against the concentration of MTs, and the data were fit to the single-site saturation non-linear regression curve (with no non-specific binding but background and *K_d_*>0 constrain) using GraphPad Prism 6.0 (San Diego, CA). The *K_d_* values generated represent the average and propagated error from at least three independent experiments.

For co-sedimentation assays with subtilisin-treated microtubules, 20 μM taxol-stabilized microtubules were incubated for 30 min at 30°C with 200 μg/ml subtilisin A (Calbiochem), a serine protease that cleaves the C-terminal 10–20 amino acids from α-and β-tubulin. The reaction was stopped with 10 mM PMSF, digested MTs were pelleted (90,000 rpm, TLA100, 10 min, 25 °C) and resuspended in the original volume of GT buffer. The assay was then carried out as described above.

Curved tubulin oligomers or rings were generated by incubating tubulin with Dolastatin (Tocris Bioscience) dissolved in dimethyl sulfoxide (DMSO) in a 2:1 molar ratio for 1 h at room temperature in GT buffer supplemented with 6% DMSO. The rings were purified by ultracentrifugation through a cushion of GT buffer with 40 % glycerol. The rings were resuspended in the original volume of GT buffer and assay was carried out as described before. Sheared microtubules were prepared by passing taxol-stabilized microtubules through a 26_1/2_-gauge needle multiple times immediately before the addition to the protein sample. The generation of increased number of ends was confirmed by transmission electron microscopy (TEM).

#### Total internal reflection fluorescence microscopy (TIR-FM)

Flow chambers containing immobilized MTs were assembled as described (McKenney et al., 2014). Imaging was performed on a Nikon Eclipse Ti-E microscope equipped with an Andor iXon EM CCD camera; a 100×, 1.49 NA objective and 1.5x tube lens (yielding a pixel size of 106 nm); four laser lines (405, 491, 568, and 647 nm); and using the Micro-Manager software (Edelstein et al., 2014). All assays were performed in assay buffer (30 mM Hepes [pH 7.4], 50 nM K-acetate, 2 mM Mg-acetate, 1 mM EGTA, and 10% glycerol), supplemented with 0.1 mg/mL biotin-BSA, 0.5% Pluronic F-168, and 0.2 mg/mL κ–casein. A final concentration of 1 and 5 nM Cdt1^92-546^ was used. Protein samples from two independent purification batches were used to ensure biological reproducibility. Photobleaching tests (Helenius et al., 2006) revealed the photobleaching rate of this construct under our conditions exceeded the characteristic dwell time by approximately three-fold and thus our dwell time calculations do not take photobleaching into account. Dwell times were calculated manually from kymographs of individual MTs and plotted in GraphPad Prism 7.0.

For measuring the diffusion coefficient, the TIR-FM imaging time series of Cdt1^92-546^ were first bleach-corrected by histogram matching using Image J Software and saved as 8-bit multi-page TIFF files before being transferred to the DiaTrack 3.04 Pro particle tracking Software (http://www.diatrack.org/), which was then used to identify and track GFP particles localized on microtubules. The read-out for diffusion coefficient obtained from the tracking Software was in (pixel)^2^/sec and the time interval between individual time frames of the TIR-FM time series was ~65 milliseconds. Thus, each read-out in (pixel)^2^/sec was first multiplied by the pixel size in μm (0.106 μm), and again by a factor of ~15.4 (milliseconds to seconds conversion) to obtain final values in (μm)^2^/sec. The particles that showed high diffusion were excluded for the calculation of the diffusion coefficient (D), along with the stationary particles. This was done because in the time frame used to conduct the analysis, we were unable to ascertain with accuracy whether these particles were indeed bound to and diffusing along MTs.

### *in vitro* phosphorylation assay

*In vitro* kinase assay was performed on 1 μg each of hekCdt1 and human Vimentin (ab73843, Abcam) at 30°C for 1 h in assay buffer containing 25 mM Hepes, 50 mM NaCl, 1 mM DTT, 2 mM EGTA, 5 mM MgSO_4_, 10 μM ATP, 5 μCi γ-[^32^P] ATP (Perkin Elmer) in the presence of 0.5 μg Aurora B Kinase (AURKB-231H, Creative BioMart) alone or with its inhibitor ZM447439 (5 or 10 μM as indicated, obtained from Sigma). Samples (30 μl) were then electrophoresed on SDS-PAG and visualized by autoradiography. The same gel was stained with coomassie to ascertain migration of the proteins.

#### Sample preparation for Mass spectroscopy, LC-MS/MS and Data analysis

Both non-radioactive kinase treated and control samples were precipitated overnight at - 20°C with 8 volumes of acetone and 1 volume of TCA. The protein was pelleted by centrifugation at 15,000 × g for 15 min at 4°C. The pellets were washed twice with 200 μL cold acetone, dried and re-suspended in 8 M urea/0.4 M ammonium bicarbonate, then reduced with 4 mM DTT for 30 min at 50°C. Alkylation with 18 mM iodoacetamide was carried out at room temperature for 30 min in dark. The urea was diluted to less than 2 M with ultrapure water followed by addition of sequencing-grade trypsin (Promega) for overnight in a ratio of 1:18 (enzyme: substrate). The peptides were desalted using C18 spin columns according to the manufacturer’s instructions. Samples were re-suspended in 5 % acetonitrile (ACN) /0.1 % formic acid (FA) to a final concentration of 0.15ug/μL.

Nano LC-MS/MS analyses were performed with a 75 μm x 10.5 cm PicoChip column packed with 3 μm Reprosil C18 beads. A 150 μm × 3 cm trap packed with 3um beads was installed in-line. Solvent A consisted of 0.1 % FA in water and solvent B was 0.1 *%* in ACN. Peptides were trapped at 5 μL/min for 5 min, then separated at a flow rate of 300 nL/min with a gradient from 5 % to 30 % B in 95 min. After a 5 min ramp to 60 % the column was washed at 95 % B and re-equilibrated to 5 % B with a total analysis time of 120 min. Desalted phosphopeptide-enriched fractions were re-suspended in 5 % ACN/0.1 % FA. Six microliters (0.9 μg) were injected in duplicates for each sample.

The LC was coupled by electrospray to a Q-Exactive HF mass spectrometer operating in data-dependent MS/MS mode with a top-15 method. Dynamic exclusion was set to 60 s and charge 1+ ions were excluded as well. MS^1^ scans were collected from 300-2000 *m/z* with resolving power equal to 60,000 at 400 *m/z.* The MS^1^ AGC was set to 3 × 10^6^. Precursors were isolated with a 2.0 m/z isolation width, and the HCD normalized collision energy was set to 35%. The MS^2^ AGC was set to 1×10^5^ with a maximum ion accumulation time of 120ms, and the resolving power was 30,000.

MS raw files were searched against the human SwissProt database (version downloaded 8/2016, 20191 sequences) using the Mascot search engine. Cysteine carbamidomethylation was a fixed modification and the variable modifications were oxidized methionine, acetylation of protein N-terminus, deamidation of Asn/Gln, and phosphorylation of Ser, Thr, and Tyr residues. Two missed cleavages were allowed. Statistical cutoffs and data visualization were accomplished using Scaffold Q+ software (Proteome Sciences). Proteins were selected with a 1 % FDR cutoff and peptides with a 90 % identification probability or better were considered. All phosphopeptide spectra were inspected manually to determine whether the phosphate group had been assigned to the correct amino acid.

### Cell culture methods

#### Transient or stable transfections, immunofluorescence and confocal microscopy

HeLa or HEK 293 cells were cultured at 37°C with 5 % CO_2_ in Dulbecco’s modified eagle’s medium (DMEM, Life Technologies) containing 10 % fetal bovine serum (Seradigm, VWR Life Science), 100 U ml^−1^ penicillin and 1 μg ml^−1^streptomycin.

Prior to transfection, cells were seeded overnight onto 18 mm circular coverslips to achieve ~ 60-70 % confluency. The cells were then transfected with 0.5 μg of pJC13-Cdt1-HASH plasmid (carrying C-terminal tags: 2× HA, Strep-tag, 12× His tag followed by a stop codon; a generous gift from Jeanette Cook at the UNC) using Effectene transfection reagent (Qiagen) according to the manufacturer’s protocol. After 36 h, the cells were fixed in phosphate buffer saline (pH 7.2) containing 4 *%* formaldehyde followed by staining with antibody against the HA tag (1:100, H6908, Sigma), Tubulin monoclonal antibody (1:500, T9026, clone DM1A Sigma) and 4,6-diamino-2-phenylindole (DAPI) dihydrochloride (1:10,000, Life Technologies). Alexa Fluor 488-, Rhodamine Red-X-, or Cy5-labeled donkey secondary antibodies used at 1:250 dilution each were obtained from Jackson ImmunoResearch Laboratories, Inc.

To generate stable cell lines, pQCXIP, a bicistronic retroviral expression vector was used (Julius et al., 2000). Upon transfection into HEK-293T packaging cell line, this vector integrates and stably expresses a viral genomic transcript containing the CMV immediate early promoter, gene of interest, IRES and the puromycin resistance gene (Pur^R^). The gene of interest and the puromycin resistance gene are co-transcribed, via the internal ribosome entry site (IRES), as a bicistronic message (Adam et al., 1991; Goshima and Yanagida, 2000). HEK-293T cells were seeded at a density of 0.75 × 10^6^ in 10-cm dish. A three plasmid transfection system containing carrier pQCXIP plasmid (5 μg) expressing the gene of interest (*cdt1*-WT or 10D or 10A) along with the two helper plasmids pVSV-G, 1.25 μg (Clonetech) and pCL-eco, 3.75 μg (Imagenex) encoding the Env and Gag/Pol proteins required for virus production was used. The HEK cells were transfected using calcium phosphate for 48 h and the culture supernatant containing viral particles was harvested and passed through 0.45 μm syringe filter. Fresh media was added to the dish for second round of collection at 72 h. Next, polybrene (8 μg/ml) was mixed with the filtered virus followed by addition of the virus on to the target HeLa cells (seeded at a density of 6.5 × 10^4^ in a 10-cm dish) for 6 h. Selection for the stably integrated cells was performed for 2 weeks by addition of 1 μg/ml puromycin, a concentration determined to kill 100 % of control un-transduced HeLa cells.

For image acquisition, the coverslips were mounted using ProLong^®^ Gold Antifade reagent (Invitrogen), 3D stacks were obtained sequentially at 200-nm steps along the z axis through the cell using a high-resolution inverted microscope (Eclipse TiE; Nikon) equipped with a spinning disk (CSU-X1; Yokogawa Corporation of America), an Andor iXon Ultra888 EMCCD camera, and an x60 or x100 1.4 NA Plan-Apochromatic DIC oil immersion objective (Nikon). The images were acquired and processed using the NIS elements Software from Nikon.

#### siRNA transfections, double thymidine synchronization and cold stability assay

HeLa cells were synchronized by treatment with 2 mM thymidine for 18 h followed by release for 9 h and then re-treatment with 2 mM thymidine for 18 h. Synthetic duplexed RNA oligonucleotide (siRNA) against Cdt1 was transfected into HeLa cells according to the manufacturer’s instructions as described previously (Varma et al., 2012) during the 2^nd^ thymidine release and prior to fixing the cells at designated time points as indicated in figure 4 and associated legends. Western blot was performed using anti-Cdt1 antibody (H-300 Santa-Cruz Biotechnology, Inc.) to evaluate the efficiency of Cdt1 knockdown. GAPDH was probed for after stripping the same Western blot as loading control. Other cell manipulations include cold treatment for 15 min with ice-cold PBS prior to fixation.

#### STLC washout assay

HeLa cells were double-thymidine synchronized and siRNA against Cdt1 was added as described above. STLC, an analog of Monastrol (7μM, Millipore Sigma) was added to the cells. After 2 h, cells were either fixed (t=0) or washed twice with and released into fresh medium containing MG132 for 1 h (t=60), before fixation and immunofluorescence staining.

#### Live Cell imaging

HeLa cell vector controls or those expressing either *wt* or Aurora B mutant versions of Cdt1 cultured on 35mm glass-bottomed dishes (MatTek Corporation) were subjected to the same thymidine and Cdt1 siRNA treatment regime described above and imaged live starting at 9 hrs after release from 2^nd^ thymidine treatment. The chromosomes were labelled with the live cell DNA dye, Hoechst (45 min, xx μg/ml), prior to the initiation of live imaging. Just before imaging, the Hoechst-containing medium was replaced with pre-warmed L-15 medium (Gibco) supplemented with 10% fetal bovine serum and fresh Cdt1 siRNA mix. Live-imaging was carried-out using an incubation chamber for microscopes (Tokai Hit Co., Ltd) at 37 ^o^C and 5% CO_2_. Images were recorded using a high-resolution inverted microscope (Eclipse TiE; Nikon) equipped with a spinning disk (CSU-X1; Yokogawa Corporation of America), an Andor iXon Ultra888 EMCCD camera and a x60 1.4 NA Plan-Apochromatic DIC oil immersion objective (Nikon) fitted with an objective heater. 4-6 1.5 μm-separated z-sections covering the entire volume of the cell were collected at every 10 min for up to 12 h. Image processing was performed using the NIS elements Software from Nikon.

#### Ni^2^-NTA agarose-mediated pull down

HEK-293T cells were transfected with the plasmids (Cdt1-HASH or cy-Cdt1 mutant, RRL (68-70) AAA which prevents Cyclin/Cdk binding, in pQCXIP and pEGFP-N1 as control) for 4 h at 20-30% confluency using PEI max transfection reagent. M phase synchronization was done by treating the cells with 2 mM thymidine for 18 h, followed by addition of 100 ng/ml (0.33 μM) Nocodazole for 10 h. The cells were collected by mitoticshake-off and lysed using lysis buffer (50 mM HEPES pH 8.0, 33 mM KAc, 117 mM NaCl, 20 mM Imidazole, 0.1% triton, 10% glycerol, 0.1 mM AEBSF, 10 μg/ml pepstatin A, 10 μg/ml aprotinin, 10 μg/ml leupeptin, 1 mM ATP, 1 mM MgCl2, 5 μg/ml phosvitin, 1 mM β-glycerol-phosphate, 1 mM orthovanadate). The whole cell lysates expressing the His-tagged proteins were incubated with Ni^+2^-NTA beads for 3 h with end-on rotation at 4°C. Beads were washed three times with the lysis buffer, and bound proteins were eluted by boiling for 5 min in 40 μL of 2× SDS sample buffer. For immunoblotting, proteins were electrophoresed on SDS-PAG and transferred to the PVDF membranes. Immunoblots were developed using chemiluminescence and exposed on to the X-ray films. Antibodies against Aurora B Kinase and HA-tag were procured from Abcam (ab2254) and Sigma (H6908), respectively.

#### Statistical analysis

GraphPad Prism software was used for determining statistical significance of the data obtained. A standard non-parameteric unpaired two-tailed/sided *t* test was used. This test assumes that both groups of data are sampled from Gaussian populations with the same standard deviation. Data distribution was assumed to be normal but this was not formally tested.

#### Online Supplemental Material

Fig. S1 (A) shows the topology of a conventional WTH domain. Fig. S1 (B and C) shows that control GST and GFP tags do not bind to MTs. Fig. S1 (D-F) shows Cdt1 binding to straight, curved and sheared MTs. Fig. S1 (G) is the TIR-FM image of Cdt1^92-546^ decorating MTs and Fig. S1 (H) demonstrates Cdt1 co-localization with MTs in HeLa cells. In Fig. S2 are the chromatograms from phosphoproteomics showing phosphorylation of full length Cdt1 (1-546 aa) by Aurora B kinase at the indicated residues. Fig. S3 (A and B) demonstrates normal mitotic spindle structure in cells rescued with Aurora B kinase phospho-mutants of Cdt1 at 37 °C and the localization of Hec1 in these cells. Video 1 shows diffusion of Cdt1 on MTs in TIR-FM experiment. Videos 2 and 3 provide Z-series data of Cdt1-HA co-localization on MTs and kinetochores, Videos 4-7 are live imaging analysis of HeLa cells rescued with Aurora B kinase phospho-mutants of Cdt1 after labeling the chromosomes with DNA dye, Hoechst.

## Supplementary Movie Legends

**Supplementary movie S1:** Continuous live TIR-FM imaging of Cdt1^92-546^ (green) and surface immobilized Dylight405-labeled MTs (blue). The total duration of the time series spanned for 129 seconds with the time interval between individual time frames as ~65 milliseconds.

**Supplementary movie S2:** Individual frames from a Z-stack obtained from spinning disc confocal microscopy of Cdt1-HA (in green) and Hec1 (kinetochore marker, in red) was assembled into a time series and played at 2 frames/sec. The individual Z-stacks were separated by 200 nm.

**Supplementary movie S3:** Individual frames from a Z-stack obtained from spinning disc confocal microscopy of Cdt1-HA (in green) and Hec1 (a kinetochore marker, in red) was assembled into a time series and played at 2 frames/sec. The individual Z-stacks were separated by 200 nm.

**Supplementary movie S4:** HeLa cells rescued with RNAi-resistant WT-Cdt1 after the knockdown of endogenous Cdt1 was subjected to live imaging after labeling the chromosomes with DNA dye, Hoechst. Z-Projections for individual time points (obtained from 4-6 1.5 μm z-sections) separated by 10 min intervals from data collected for a total period of 170 min (for this example) were sped up to ~ 28× and played at ~2.8 frames/sec.

**Supplementary movie S5:** HeLa cells rescued with the control plasmid after the knockdown of endogenous Cdt1 (simulates Cdt1-depleted state) was subjected to live imaging after labeling the chromosomes with DNA dye, Hoechst. Z-Projections for individual time points (obtained from 4-6 1.5 μm z-sections) separated by 10 min intervals from data collected for a total period of 440 min (for this example) were sped up to ~ 28× and played at ~2.8 frames/sec.

**Supplementary movie S6:** HeLa cells, rescued with RNAi-resistant Aurora B Kinase phosphomimetic Cdt1 mutant (Cdt1-10D) after the knockdown of endogenous Cdt1, were subjected to live imaging after labeling the chromosomes with DNA dye, Hoechst. Z-Projections for individual time points (obtained from 4-6 1.5 μm z-sections) separated by 10 min intervals from data collected for a total period of 450 min (for this example) were sped up to ~ 28× and played at ~2.8 frames/sec.

**Supplementary movie S7:** HeLa cells, rescued with RNAi-resistant Aurora B Kinase non-phosphorylatable Cdt1 mutant (Cdt1-10A) after the knockdown of endogenous Cdt1, were subjected to live imaging after labeling the chromosomes with DNA dye, Hoechst. Z-Projections for individual time points (obtained from 4-6 1.5 μm z-sections) separated by 10 min intervals from data collected for a total period of 170 min (for this example) were sped up to ~ 28× and played at ~2.8 frames/sec.

**Supplementary figure 1.**
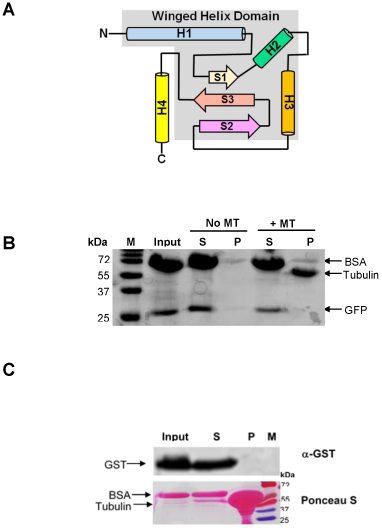
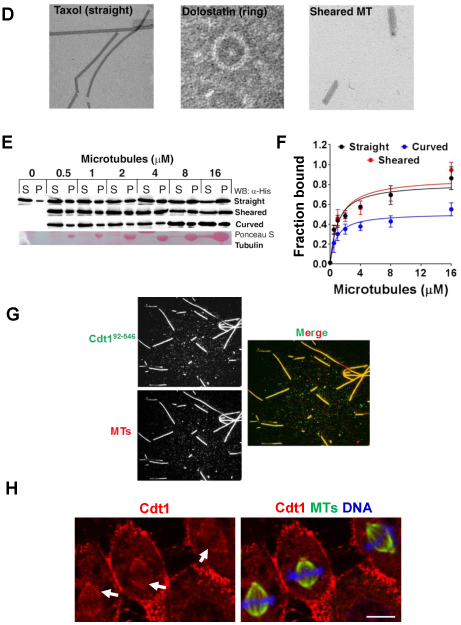
**(A)** Topology diagram showing the elements of the canonical winged-helix domain highlighted in the grey box; H and S depict α-helix and β-sheets, respectively. **(B)** Coomsassie stained gel of GFP (2 μM) alone in supernatant and pellet pre and post incubation with microtubules (4 μM). The total amount of input protein used in the assay after pre-clarification is also indicated. M represents migration of molecular mass standards on an 18% SDS-PAG. **(C)** Western blot with anti-GST antibody depicting partitioning of GST (2 μM) alone in the supernatant and pellet fraction in the presence of microtubules (4 μM). The total amount of input protein used in the assay after pre-clarification is also indicated. M represents migration of molecular mass standards on an 18% SDS-PAG. The lower panel depicts the same blot stained with Ponceau S to detect the presence of tubulin in the pellet fraction. **(D)** Electron micrographs of negatively stained taxol-stabilized straight, dolostatin-treated ring/curved or sheared MTs. **(E)** Representative western blots of Cdt1^92-546^ interaction with MTs which are differentially treated/processed; probed with anti-His antibody. **(F)** Quantification of Cdt1^92-546^ (1 μM) bound to MT in a co-sedimentation assay with the indicated MT type and concentrations using image J. The same blots were stained with Ponceau S after developing with chemiluminescence. **(G)** Single channel and merged TIR-FM images showing surface immobilized Dylight405-labeled MTs (pseudo-colored in red) and 100 nM Cdt1^92-546^ protein (in green). **(H)** A low magnification representative image showing multiple HeLa cells with endogenous Cdt1 staining co-localized with spindle MTs (indicated by arrows). Red is Cdt1 antibody marking endogenous Cdt1, green is the Tubulin antibody staining MTs and blue is DAPI staining the chromosomes. Scale bar represents 10 μm.

**Supplementary figure 2.**
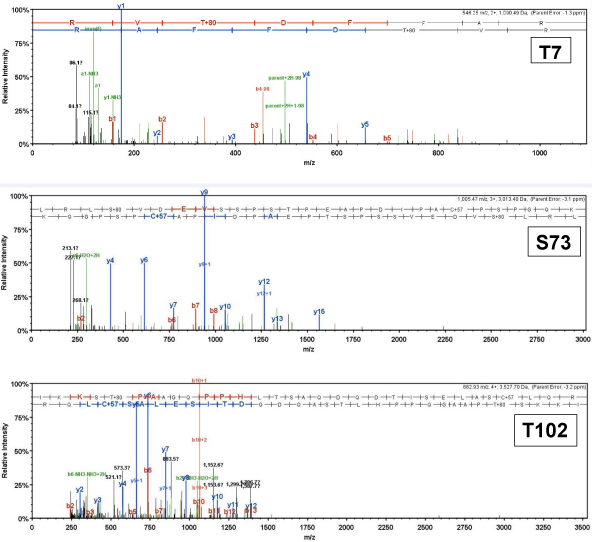
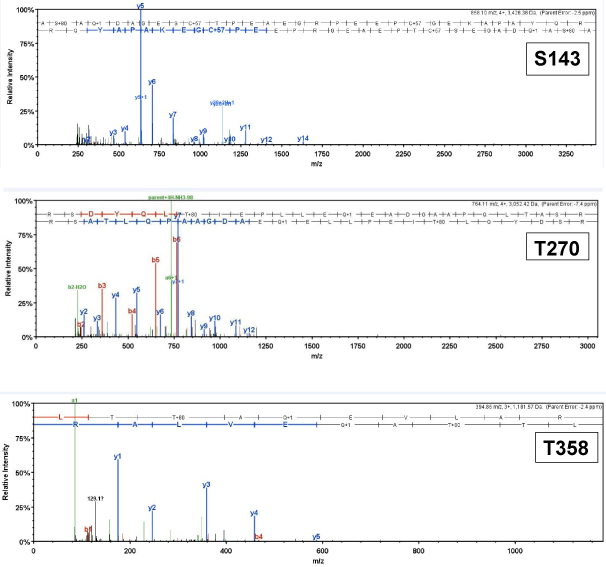
Chromatograms from phosphoproteomics showing phosphorylation of full length Cdt1 (1-546 aa, obtained from Origene with C-terminal myc/DDKK tag) at particular residues indicated on the top right hand side in a box, exclusively upon Aurora B Kinase addition (T7, T102, S143, and T358) or as inherently phosphorylated (S73 and T270) even in the kinase untreated sample indicating at least a potential phosphosite.

**Supplementary Figure 3.**
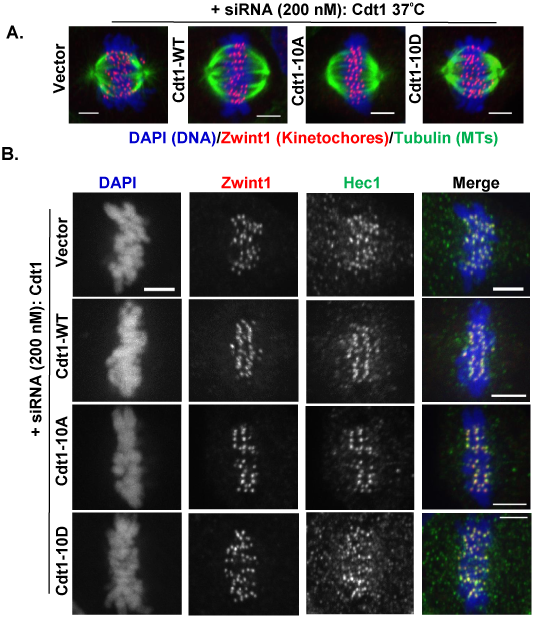
**(A)** Synchronized HeLa cells treated with siRNA against endogenous Cdt1 rescued with stably expressing siRNA-resistant Cdt1-WT or 10D (phosphomimetic) and 10A (phosphodefective) proteins were stained with antibodies against Zwint1 in red (1:500 dilution, kinetochore marker), microtubules in green (1:500 dilution, spindle marker) and DAPI in blue marks the chromosomes. Control cells that underwent extended incubation at 37°C before being fixed and stained in parallel to those exposed to cold. **(B)** Synchronized HeLa cells treated with siRNA against endogenous Cdt1 rescued with stably expressing siRNA-resistant Cdt1-WT or 10D (phosphomimetic) and 10A (phosphodefective) proteins were stained with antibodies against Hec1 in green (1:500 dilution), Zwint1 in red (1:450 dilution) (as a kinetochore marker) and DAPI in blue marks the chromosomes.

